# Density-Dependent Rearrangement of Desmoplakin in Epithelial Junctions: Insights from Expansion Microscopy

**DOI:** 10.64898/2026.04.30.721849

**Authors:** Ukyo Kawasaki, Yusuke Arakawa, Tetsuhisa Otani, Kazumasa Ohashi, Shuhei Chiba

## Abstract

Desmosomes mechanically couple neighboring epithelial cells to the intermediate filament cytoskeleton. However, how the nanoscale organization changes with epithelial cell density remains incompletely understood. Here, we used expansion microscopy (ExM) to examine Madin–Darby canine kidney epithelial monolayers. ExM revealed density-associated differences in the relative apicobasal organization of epithelial junctions, with increased separation between occludin-positive tight junctions and desmoglein 2-positive desmosomes in confluent monolayers. A comparison of subconfluent and confluent cultures revealed a density-associated reorganization of desmoplakin (DSP) within opposing desmosomal plaques, with a more pronounced change in C-terminal spacing than in N-terminal spacing. Independent STED analysis of non-expanded samples confirmed the density-associated increase in DSP C-terminal spacing. In keratin filament-deficient cells, this spacing was elevated in subconfluent cultures and did not increase with cell density. Pharmacological inhibition of myosin II or ROCK reduced this elevated spacing, indicating that keratin loss-associated reorganization of DSP is sensitive to actomyosin contractility. These findings show that epithelial cell density is associated with the remodeling of epithelial junction organization and DSP plaque architecture, demonstrating the utility of ExM for quantitative analysis of junctional architecture.

**Summary statement:** Expansion microscopy reveals how epithelial cell density and the cytoskeleton are linked to nanoscale remodeling of desmosomes, the junctions that mechanically connect neighboring cells.

## Introduction

Epithelial tissues rely on specialized intercellular junctions to maintain their structural integrity. Along the apicobasal axis, tight junctions (TJs) form an apical belt above adherens junctions (AJs) and desmosomes (Farquhar and Palade, 1963; Janssen and Huveneers, 2024). Desmosomes are intermediate filament (IF)-anchored cell–cell junctions that support epithelial mechanical integrity (Harmon and Green, 2013; Yeruva and Waschke, 2023). Electron microscopy established their symmetric plaque architecture, comprising an outer dense plaque (ODP) adjacent to the plasma membrane and an inner dense plaque (IDP) associated with IFs (Farquhar and Palade, 1963; North et al., 1999; Bharathan et al., 2024).

Super-resolution microscopy and cryo-electron tomography have refined this architecture at the molecular scale (Bharathan et al., 2024; Stahley et al., 2016). Current models place the cytoplasmic tails of desmogleins (DSGs), desmocollins (DSCs), plakoglobin (PG), and plakophilins (PKPs) in the ODP, whereas desmoplakin (DSP) extends from the ODP to the IDP to link IFs (Fig. 1A). Desmosomal plaques exhibit structural plasticity, with previous studies showing remodeling associated with the adhesive state, epidermal differentiation, and junction assembly (Beggs et al., 2022; Stahley et al., 2016). Structural changes have also been reported in pemphigus (Schmitt et al., 2025). Actomyosin-dependent mechanical inputs have been implicated in desmosome regulation, and recent biophysical evidence supports mechanical coupling among keratin filaments, actin, and DSP (Moch et al., 2022; Dong et al., 2025).

**Figure 1.**
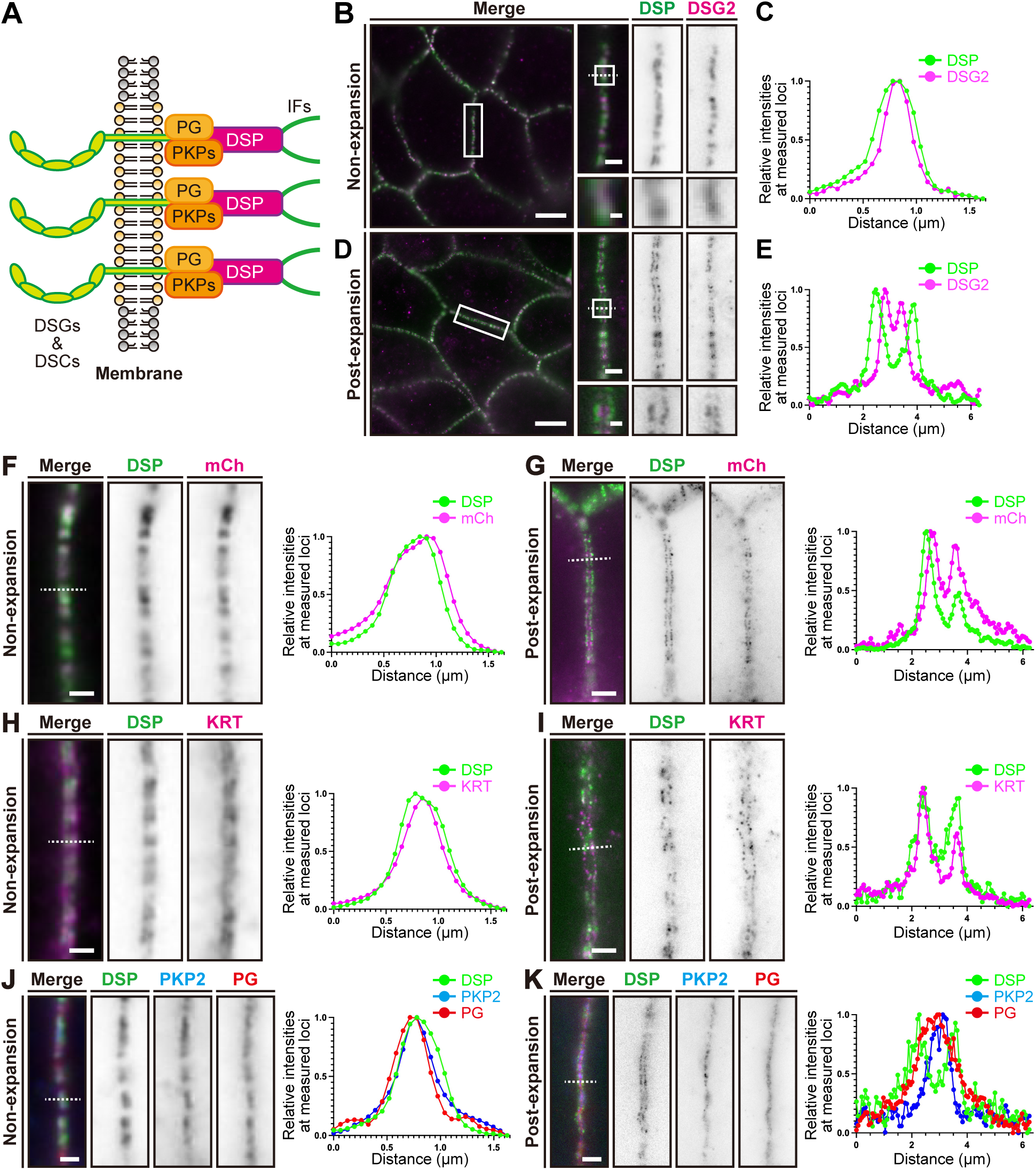
Plaque-scale desmosome organization resolved by ExM. (A) Schematic of desmosome architecture. PG, plakoglobin; PKPs, plakophilins; DSP, desmoplakin; DSGs, desmogleins; DSCs, desmocollins; IFs, intermediate filaments. (B– E) Representative non-expanded (B) and expanded (D) confluent MDCK cells stained for DSP and DSG2, with corresponding line-scan profiles (C, E). (F–K) Representative non-expanded (F, H, J) and expanded (G, I, K) confluent MDCK cells stained for mSca3-DSP (mCherry) and DSP C-terminus (F, G), DSP and pan-keratin (KRT; H, I), or DSP, PKP2, and PG (J, K), with corresponding line-scan profiles. Scale bars: 5, 1 and 0.25 µm (B) and 20, 4 and 1 µm (D), from overview to successive magnifications; 1 µm (F, H, J); 4 µm (G, I, K).

Taken together, these studies indicate that desmosomal organization is remodeled in both physiological and pathological contexts. However, how epithelial cell density relates to the relative apicobasal organization of epithelial junctions and desmosomal plaque organization, particularly the contribution of the keratin cytoskeleton, remains incompletely understood. Therefore, we used expansion microscopy (ExM) (Chen et al., 2015; Tillberg et al., 2016) to quantitatively analyze these features in subconfluent and confluent Madin–Darby canine kidney (MDCK) monolayers and examined the contributions of keratin filaments and actomyosin contractility.

## Results and Discussion

### Plaque-scale desmosome organization resolved by ExM

Conventional fluorescence microscopy readily visualizes overall desmosome distribution and assembly, but diffraction-limited resolution restricts quantitative analysis of internal plaque-scale organization (Fig. 1B, C). To resolve opposing plaque signals in confluent MDCK monolayers, we applied the Uni-ExM protocol (Cui et al., 2023), which physically expands biological samples by approximately 4–4.5× for sub-diffraction imaging (Chen et al., 2015; Tillberg et al., 2016). At each desmosome, cytoplasmic plaques from the two adjacent cells face each other across the cell–cell interface. In non-expanded samples, the DSP C-terminal signals from these opposing plaques largely overlapped and could not be consistently resolved as separate peaks. After expansion, these signals were spatially separated and resolved into two distinct peaks (Fig. 1B–E). The desmoglein 2 (DSG2) C-terminal signal was also resolved into two peaks between the DSP peaks, consistent with its membrane-proximal position (Stahley et al., 2016; Bharathan et al., 2024).

We next characterized expansion across the spatial scales relevant to these measurements. The macroscopic post-/pre-expansion gel diameter ratio was 4.2 ± 0.1 (Fig. S1A, B). In same-cell analyses, post-/pre-expansion ratios of relative vector lengths and included angles were 1.00 ± 0.06 and 1.01 ± 0.07, respectively, indicating largely preserved in-plane geometry (Fig. S1C, D). At the desmosomal scale, comparison of measurements obtained from ExM and stimulated emission depletion (STED) microscopy images yielded ratios of 4.29 ± 0.42 for DSP plaque-to-plaque distance and 4.29 ± 0.93 for plaque length (Fig. S1E–H). Thus, desmosomal-scale measurements were consistent with an approximately 4.2-fold macroscopic expansion. Based on a lateral optical resolution of approximately 250–300 nm, the estimated effective lateral resolution after expansion was approximately 60–70 nm.

To map the DSP organization within the plaque, we used N-terminally mScarlet3-tagged DSP (mSca3-DSP), which restored junctional DSG2 and plakophilin 2 (PKP2) accumulation and keratin association in *DSP*-knockout (*DSP-*KO) cells (Fig. S2). ExM revealed distinct N- and C-terminal signal distributions (Fig. 1F, G), consistent with previous direct stochastic optical reconstruction microscopy (dSTORM) data (Stahley et al., 2016). The DSP C-terminal signal was adjacent to keratin filaments (Fig. 1H, I), consistent with its established keratin association (Moch et al., 2020; Stappenbeck et al., 1993).

Next, we examined PKP2 and PG. Both proteins were detected as single compact fluorescence peaks across the junction interface, and their opposing signals were not resolved separately (Fig. 1J, K). In contrast, opposing DSG2 C-terminal signals were resolved as two peaks under our imaging conditions. This difference may reflect a larger effective separation of the antibody-recognized DSG2 epitopes across the two apposed plaques, although differences in antibody labeling and epitope accessibility may also contribute (Stahley et al., 2016; Bharathan et al., 2024). Together, these observations demonstrated the utility of ExM in characterizing plaque-scale desmosomal organization and its spatial relationship with the keratin network in confluent MDCK monolayers.

### Apicobasal organization of epithelial junctions differs with cell density

To compare the relative apicobasal positions of epithelial junctions, we examined the axial distribution of occludin (OCLN), β-catenin, and DSG2 in subconfluent and confluent MDCK monolayers (Fig. 2). Substrate coverage and cell area confirmed distinct epithelial density states (Fig. S3A–C). In non-expanded samples, OCLN, β-catenin, and DSG2 signals were observed in close proximity in subconfluent cells, whereas in confluent cells, OCLN and DSG2 were more apically localized and β-catenin was broadly distributed along the lateral membrane (Fig. 2A–G). After expansion, the relative axial positions of OCLN and DSG2 were more clearly resolved in subconfluent cells (compare Fig. 2B, C with I, J), whereas in confluent cells the OCLN– DSG2 separation was more pronounced, with DSG2 positioned basal to OCLN (compare Fig. 2E, F, G with L, M, N). β-catenin showed a broader axial distribution distinct from both OCLN and DSG2. In confluent cultures, the DSG2 peak was approximately 1.8 µm basal to OCLN in expanded samples, corresponding to approximately 430 nm in the native dimension and consistent with previous ultrastructural observations (Rajasekaran et al., 1996). Taken together, these results showed that the relative apicobasal junction organization differed with epithelial cell density, with more pronounced OCLN–DSG2 separation in confluent MDCK monolayers.

**Figure 2.**
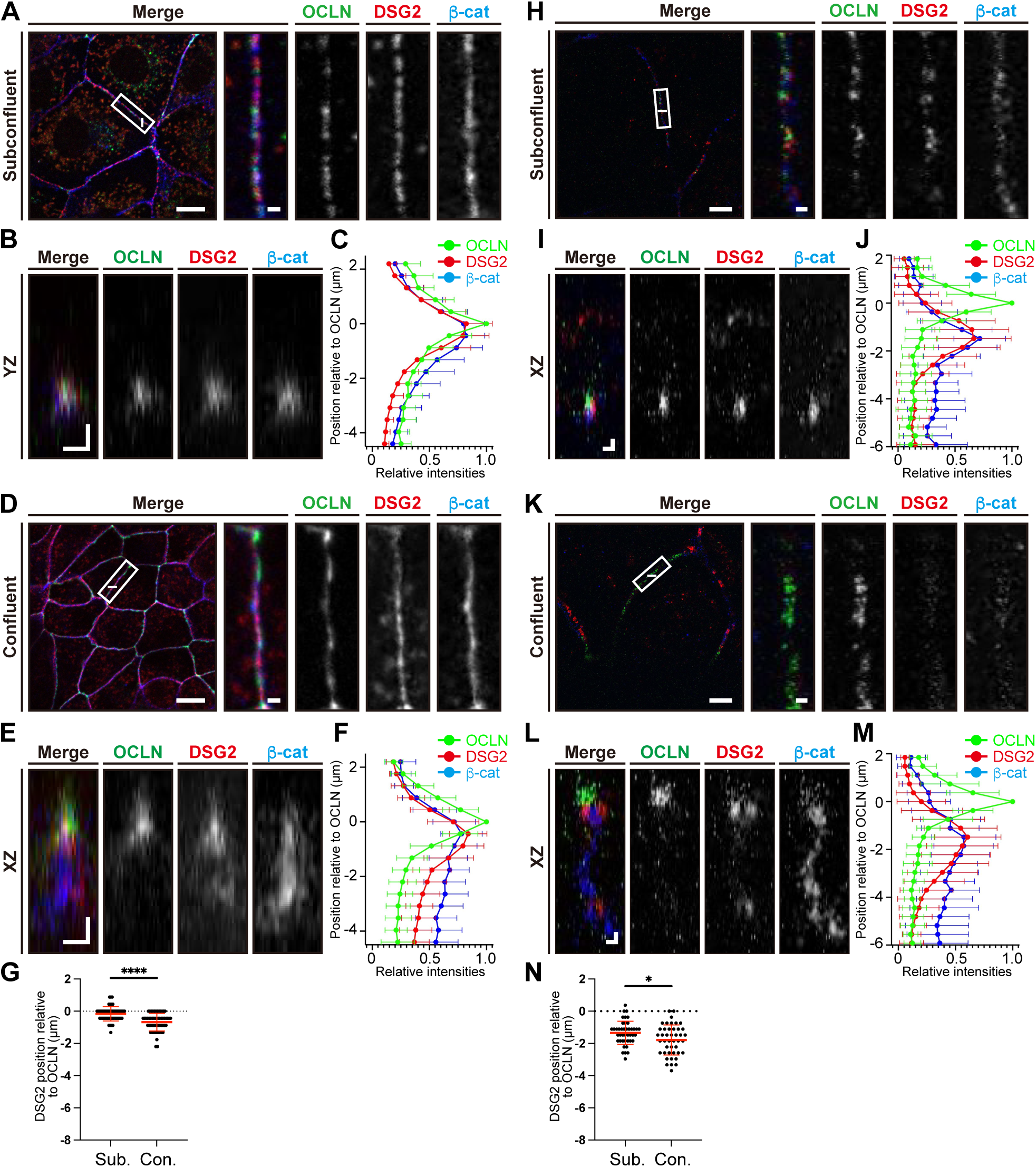
Apicobasal organization of epithelial junctions resolved by ExM. (A–G) Non-expanded subconfluent (A–C) and confluent (D–F) MDCK cells stained for OCLN, DSG2, and β-catenin (β-cat), showing representative XY single optical sections (A, D), orthogonal YZ (B) and XZ (E) views from the corresponding z-stacks at the lines indicated in A and D, mean z-axis fluorescence intensity profiles (C, F), and OCLN– DSG2 peak separation (G). For the profiles, the OCLN fluorescence maximum was defined as z = 0; positive and negative values indicate apical and basal positions, respectively. (H–N) Corresponding analysis of expanded subconfluent (H–J) and confluent (K–M) cells, showing representative XY single optical sections (H, K), XZ views (I, L), z-axis fluorescence profiles (J, M), and OCLN–DSG2 peak separation (N). Mean profiles in C, F, J, and M were averaged across 52, 61, 43, and 43 independent cell– cell junctions, respectively; error bars indicate s.d. In G and N, the horizontal lines and error bars indicate mean and s.d., respectively. Scale bars: 10 and 1 µm (A, D, H, K), from overview to magnification; 1 µm (B, E, I, L).

### DSP undergoes density-dependent reorganization

To examine how DSP plaque organization changes with epithelial cell density, we first quantified the plaque-to-plaque spacing of the DSP C-terminal signal in subconfluent and confluent cells. DSP C-terminal spacing increased from 934.2 ± 132.9 nm in subconfluent cells to 1104.7 ± 129.5 nm in confluent cells, corresponding to an increase of 170.5 nm (Fig. 3A–C). We next examined the DSP N-terminal signal using mSca3-DSP. N-terminal spacing also increased with cell density, from 711.8 ± 139.1 nm in subconfluent cells to 774.1 ± 134.9 nm in confluent cells, corresponding to an increase of 62.3 nm (Fig. 3D–F). Thus, the density-associated increase in plaque-to-plaque spacing was larger at the DSP C-terminus than at the N-terminus.

**Figure 3.**
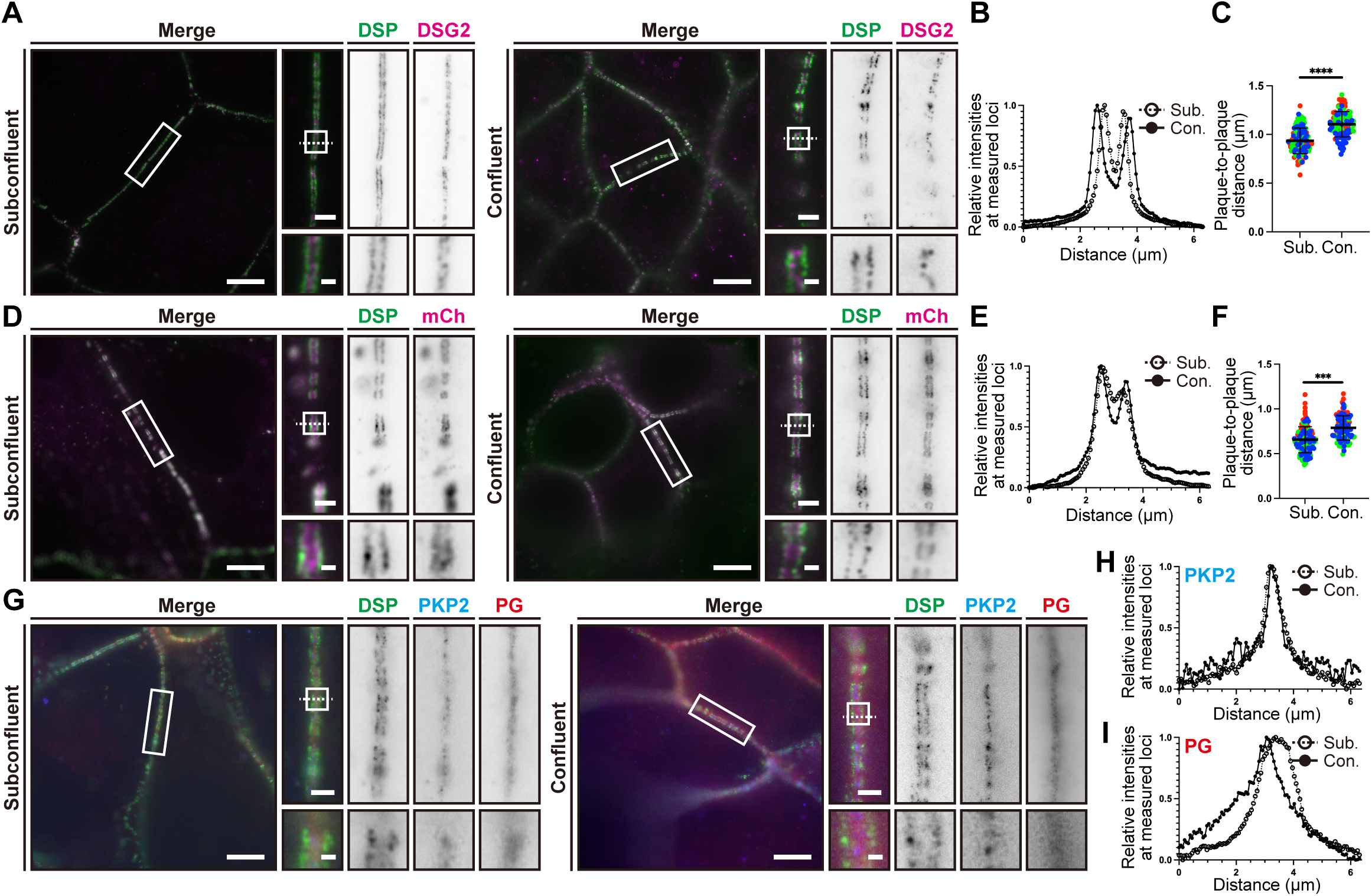
Density-dependent reorganization of DSP. (A–I) ExM analysis of subconfluent (Sub.) and confluent (Con.) MDCK cells. (A–C) Representative images of DSP C-terminus and DSG2 (A) with DSP C-terminal line-scan profiles (B) and plaque-to-plaque distance (C). (D–F) Representative mSca3-DSP-expressing cells stained for DSP C-terminus and mCherry (DSP N-terminus) (D), with DSP N-terminal line-scan profiles (E) and plaque-to-plaque distance (F). (G–I) Representative images of DSP, PKP2, and PG (G) with PKP2 (H) and PG (I) line scans. Line scans in B, E, H and I were obtained along the dotted lines. Plaque-to-plaque distances are shown in post-expansion dimensions. For C and F, each dot represents one desmosome; at least 30 desmosomes per condition were analyzed in each of three independent biological replicates. Colors indicate biological replicates; horizontal lines and error bars indicate the mean and s.d., respectively. Scale bars: 20, 4 and 1 µm, from overview to successive magnifications.

Assuming symmetric organization of opposing plaques, we estimated the intraplaque axial separation between the DSP N- and C-terminal signals as half the difference between their respective plaque-to-plaque distances. This estimated separation increased from 111.2 nm in subconfluent cells to 165.3 nm in confluent cells, corresponding to an increase of 54.1 nm. These measurements indicate that density-associated DSP reorganization is spatially non-uniform within the desmosomal plaque. Given the established localization of the DSP N-terminus within the ODP and its C-terminal region toward the IDP–keratin interface, the larger change at the C-terminus is consistent with more pronounced remodeling toward the keratin-associated region of the plaque (Stahley et al., 2016; Bharathan et al., 2024; Moch et al., 2020). The opposing PG and PKP2 signals were not resolved separately under either density condition, precluding a quantitative comparison of their plaque-to-plaque distances (Fig. 3G–I).

The density-associated increase in DSP C-terminal spacing observed here contrasts with the decreases reported during PKP1-induced hyperadhesion, epidermal differentiation, or calcium-dependent junction assembly (Stahley et al., 2016; Beggs et al., 2022). The density-associated increase was reproduced using an alternative fixation protocol and the DSP antibody employed by Beggs et al. (2022) and was independently confirmed in non-expanded samples using STED microscopy (Fig. S4A–F). In a calcium-switch assay similar to that of Beggs et al. (2022), the DSP C-terminal spacing was greater at 12 h than at 3 h after calcium restoration in both expanded samples analyzed by ExM and non-expanded samples analyzed by Airyscan 2 SR imaging (Fig. S4G–J). Together, these controls argue against fixation, antibody choice, or the ExM procedure as the sole explanations for the opposing structural responses.

### Keratin filaments regulate density-dependent DSP reorganization

Given the more pronounced density-associated change toward the keratin-associated DSP C-terminal region, we examined the contribution of keratin filaments. Because MDCK cells express multiple keratin isoforms with substantial functional redundancy (Hagiyama et al., 2017; Coulombe and Omary, 2002), we generated keratin filament–deficient (*KRT-*null) MDCK cells using a keratin gene cluster deletion strategy and confirmed the loss of keratin proteins and filamentous structures (Fig. S5A–E). In wild-type (WT) cells, DSP C-terminal spacing increased from 899.0 ± 126.4 nm in subconfluent cultures to 1174.4 ± 166.9 nm in confluent cultures. In *KRT-*null cells, C-terminal spacing was already larger in subconfluent cells than in subconfluent WT cells (1097.7 ± 144.1 nm) and did not increase further with cell density (1076.6 ± 120.8 nm in confluent cultures) (Fig. 4A, B). DSP N-terminal spacing remained comparable between subconfluent and confluent *KRT-*null cells (Fig. 4C–E). This pattern indicated that the density-associated C-terminal spacing response observed in WT cells depended on the keratin filament network (Fig. S5F).

**Figure 4.**
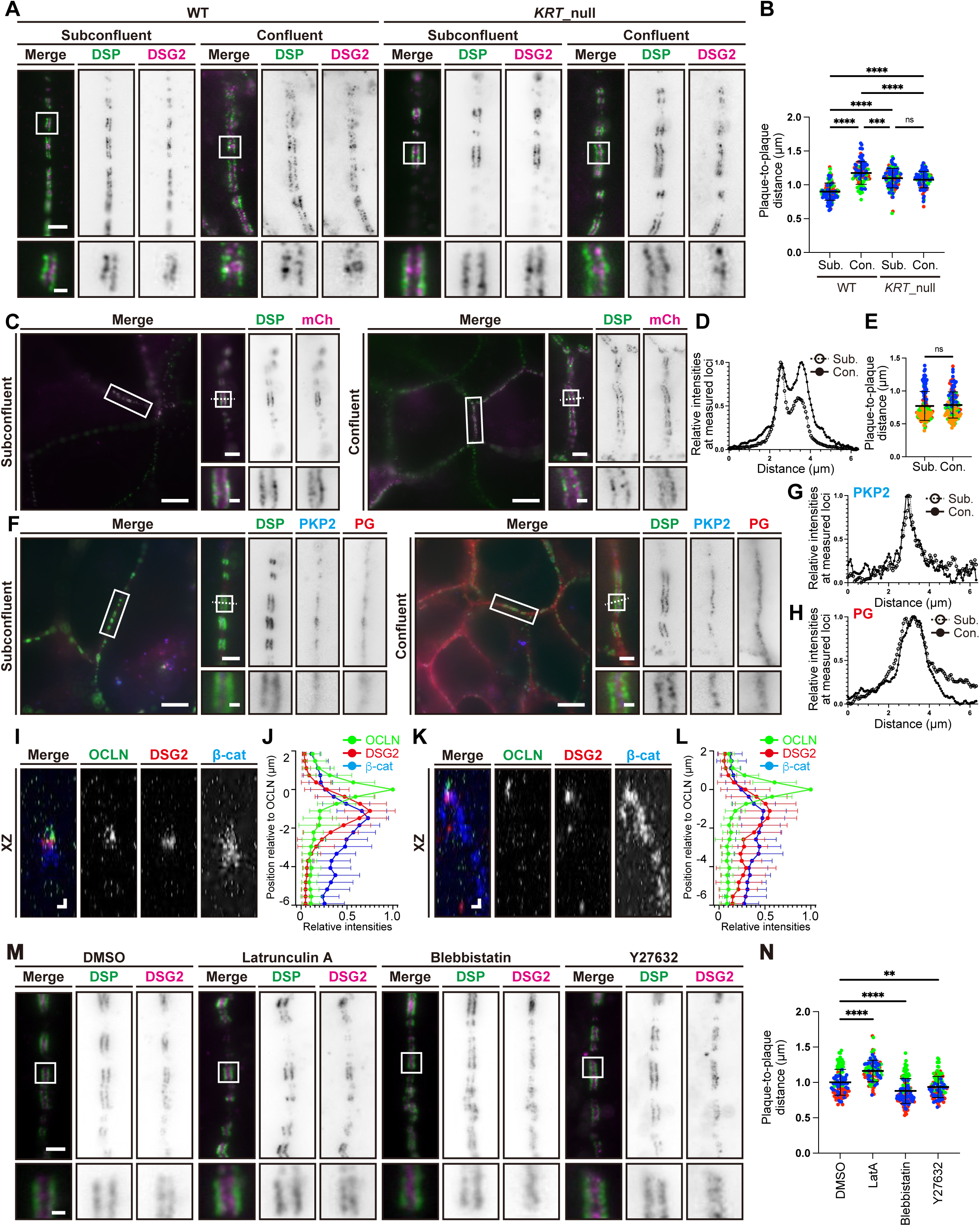
Keratin filaments regulate density-dependent DSP reorganization. (A, B) Representative ExM images of subconfluent (Sub.) and confluent (Con.) WT and *KRT-*null MDCK cells stained for DSP C-terminus and DSG2 (A), with DSP C-terminal plaque-to-plaque distance (B). (C–E) Subconfluent and confluent *KRT-*null cells expressing mSca3-DSP analyzed for DSP N-terminal spacing as in Fig. 3D–F. (F–H) Subconfluent and confluent *KRT-*null cells analyzed for PKP2 and PG as in Fig. 3G–I. (I–L) ExM analysis of subconfluent (I, J) and confluent (K, L) *KRT-*null cells stained for OCLN, DSG2, and β-catenin (β-cat), showing representative XZ views (I, K) and mean z-axis fluorescence intensity profiles (J, L). Mean profiles in J and L were averaged across 38 and 38 independent cell–cell junctions, respectively; error bars indicate s.d. (M, N) Representative ExM images of subconfluent *KRT-*null cells treated for 3 h with DMSO, latrunculin A (LatA; 0.2 µM), blebbistatin (50 µM), or Y-27632 (10 µM) before fixation, stained for DSP C-terminus and DSG2 (M), with DSP C-terminal plaque-to-plaque distance (N). Line scans in D, G and H were obtained along the dotted lines. Plaque-to-plaque distances are shown in post-expansion dimensions. For B, E and N, each dot represents one desmosome; at least 30 desmosomes per condition were analyzed in each of at least three independent biological replicates. Colors indicate biological replicates; horizontal lines and error bars indicate the mean and s.d., respectively. Scale bars: 4 and 1 µm (A, M), from overview to magnification; 20, 4 and 1 µm (C, F), from overview to successive magnifications; 1 µm (I, K).

Opposing PG and PKP2 signals were not separately resolved in *KRT-*null cells under either density condition, precluding quantitative comparison of their plaque-to-plaque distances (Fig. 4F–H). In contrast, OCLN–DSG2 apicobasal separation increased from subconfluent to confluent *KRT-*null cultures (Fig. 4I–L). This finding indicated that the loss of the DSP C-terminal density response does not simply reflect a general loss of density-associated junctional reorganization.

As DSP C-terminal spacing was already elevated in subconfluent *KRT-*null cells, we investigated whether this altered plaque organization remained responsive to actomyosin perturbation. Inhibition of myosin II with blebbistatin or Rho-associated kinase (ROCK) with Y-27632 reduced the elevated spacing, whereas disruption of F-actin with latrunculin A further increased the spacing (Fig. 4M, N). Thus, the DSP organization associated with keratin loss remains sensitive to actomyosin perturbation, with opposing responses consistent with the distinct effects of myosin-dependent contractility and F-actin integrity.

Recent biophysical evidence provides a useful framework for interpreting these responses. Dong et al. (2025) showed in MCF7 cells that actomyosin-generated forces are mechanically coupled to DSP through KRT19. In MDCK cells, they showed that actomyosin contractility shifted desmosome organization, with increased contractility producing a more extended configuration and blebbistatin producing a more compact organization. This response is directionally consistent with our observation that blebbistatin and Y-27632 reduced DSP C-terminal spacing in *KRT*-null MDCK cells. However, selective *KRT19* deletion in MCF7 cells reduced DSP C-terminal spacing without significantly altering K8 or K18 levels. In contrast, DSP C-terminal spacing was increased in our *KRT*-null cells, in which keratin gene cluster deletion broadly disrupted keratin filament formation. Therefore, these opposing responses may reflect the difference between the loss of a specific keratin isoform and the broader disruption of the keratin filament network.

During calcium-dependent junction assembly, our calcium-switch analysis showed an increase in DSP C-terminal spacing from 3 to 12 h after calcium restoration, in contrast to the decrease reported by Beggs et al. (2022). Thus, this discrepancy is unlikely to arise simply from the difference between density-associated remodeling and calcium-dependent assembly. Our MDCK cells exhibited an MDCK II-like barrier phenotype (transepithelial electrical resistance [TER], 113 ± 11 Ω·cm²; Fig. S4K; Barker and Simmons, 1981; Furuse et al., 2001; Tokuda and Furuse, 2015), whereas the MDCK subline used by Beggs et al. (2022) was not specified. Although TER alone does not establish MDCK subtype, differences in cellular background may partly account for the divergent DSP spacing responses.

Together, our findings identify a spatially non-uniform reorganization of the DSP within the desmosomal plaque during density-associated epithelial remodeling, with a larger change in the C-terminal spacing than in the N-terminal spacing. The density-associated C-terminal spacing response depends on the keratin filament network, and altered DSP organization in *KRT-*null cells remains sensitive to perturbation of myosin II, ROCK, and F-actin. The structural basis of these spacing changes remains unresolved and may involve multiple factors, including DSP orientation, plaque geometry, intercellular spacing, and intramolecular extension. Nevertheless, these findings link epithelial cell density, keratin filaments, and the actomyosin cytoskeleton to the regulation of DSP plaque organization.

## Materials and Methods

### Antibodies and expression plasmid

The antibodies used in this study are listed in Supplementary Table S1. To create the human DSP constructs, cDNA (isoform 1; NCBI RefSeq accession no. NM_004415.4) encoding a 2871-amino-acid protein was amplified by PCR using cDNA prepared from HEK293T cells. The PCR product was cloned directly into pmScarlet3_C1 (Addgene plasmid #189753; RRID: Addgene_189753; deposited by Dorus Gadella), and its sequence was confirmed.

### Cell culture and transfection

MDCK cells were obtained from the RIKEN BioResource Research Center (Japan) and maintained in Dulbecco’s Modified Eagle Medium (DMEM) containing 10% fetal bovine serum (FBS) at 37°C in 5% CO_2_. Plasmid transfection was performed using Lipofectamine LTX and Plus Reagent (Life Technologies, Carlsbad, CA, USA) as previously described (Ninomiya et al., 2024). For subconfluent and confluent cultures used for ExM and cell-density characterization, 1.5 × 10^5^ and 7.5 × 10^5^ cells, respectively, were seeded onto coverslips in 12-well plates. For non-expanded samples, 3.75 × 10^5^ and 1.86 × 10^6^ cells, respectively, were seeded onto high-precision #1.5H coverslips in 35-mm dishes. Cells were cultured for 48 h, and high-density cultures were fully confluent during this period. For pharmacological perturbation of the actomyosin cytoskeleton, subconfluent *KRT-*null MDCK cells were treated for 3 h with dimethyl sulfoxide (DMSO), latrunculin A (0.2 µM), blebbistatin (50 µM), or Y-27632 (10 µM) before fixation. *DSP*-KO cells were transiently transfected with mSca3-DSP and analyzed 48 h post-transfection. The cells were tested for mycoplasma contamination and confirmed to be negative.

### Conventional immunostaining

For non-expanded fluorescence imaging, MDCK cells were grown on 18 × 18 mm high-precision #1.5H coverslips (0.170 ± 0.005 mm thickness; Carl Zeiss, Oberkochen, Germany). Cells were fixed with ice-cold methanol for 10 min at −20°C before immunostaining. For the fixation-control experiment shown in Fig. S4A, B, parallel samples were fixed either with ice-cold methanol for 10 min at −20°C or with 50% methanol/50% acetone for 10 min at 4°C. Cells were then blocked with 2% (v/v) FBS in PBS for 30 min at room temperature (20–25°C) and stained overnight at 4°C with primary antibodies diluted with 2% (v/v) FBS in PBS or Can-Get Signal immunostaining solution for anti-pan-keratin antibodies (Toyobo, Osaka, Japan). The cells were washed three times with PBS and incubated with secondary antibodies for 1 h at room temperature. For non-expanded imaging, coverslips were mounted on glass slides using Super Clear Mount (refractive index, 1.52).

### Expansion microscopy

Expansion microscopy samples were prepared according to the Uni-ExM protocol (Cui et al., 2023) with slight modifications. Immunostained cells on coverslips were pre-incubated twice with 100 mM NaHCO_3_ (pH 8.5) for 15 min each and then incubated with anchoring buffer containing 0.04% (w/v) glycidyl methacrylate and 100 mM NaHCO_3_ (pH 8.5) overnight at room temperature. For gelation, a monomer solution (MS) containing 8.6% (w/v) sodium acrylate, 2.5% (w/v) acrylamide, 0.15% *N,N*′*-* methylenebisacrylamide, 2 M NaCl, and 1× PBS was prepared. Sodium acrylate was prepared from acrylic acid (Sigma-Aldrich, St. Louis, MO, USA) neutralized with NaOH to pH 7.5–7.8, as described previously. The gelation solution (GS) was prepared by mixing MS, 10% ammonium persulfate (APS), *N,N,N*′*,N*′-tetramethylethylenediamine (TEMED; cat. 205-06313; FUJIFILM, Tokyo, Japan) and distilled H_2_O at a 470:10:1:19 ratio. Coverslips were placed on the GS in a petri dish and incubated on ice for 15 min, followed by incubation for 60 min at 37°C to allow for gel polymerization. The coverslip-attached gels were then transferred to digestion buffer containing 50 mM Tris-HCl, pH 8.0, 1 mM EDTA, 0.5% Triton X-100, 0.8 M guanidine HCl, and 6 U/mL proteinase K for 30–45 min at 37°C. During digestion, the gels detached from the coverslips. The gels were subsequently rinsed twice with PBS for 30 min, followed by expansion with water (5–30 min) until the gel size reached a plateau. Water containing 4′,6-diamidino-2-phenylindole (DAPI) was used in the second exchange for nuclear staining. The fully expanded hydrogels were transferred to #1.5 glass-bottom dishes (coverglass thickness, 0.16–0.19 mm; Matsunami, Osaka, Japan), with the cell-containing surface facing the glass bottom and the basal surface closest to the objective. The gels were surrounded with 2% agarose to prevent drift and dehydration during imaging.

### Establishment of *DSP-*KO and *KRT-*null MDCK cell lines

To investigate the role of keratin filaments in desmosome organization, we identified the major keratin isoforms expressed in MDCK cells. RNA-seq data indicate that MDCK cells express *KRT8*, *KRT18*, *KRT7*, and *KRT19* as the predominant keratins (Hagiyama et al., 2017). Given the significant functional redundancy within the keratin gene family, which often complicates single-gene knockout studies (Coulombe and Omary, 2002), we employed a cluster deletion strategy. We identified a type-II keratin gene cluster on canine chromosome 27, which contains multiple type-II keratin genes along with the type-I keratin gene *KRT18* (Herrmann and Aebi, 2016; Vijayaraj et al., 2009). To delete this entire cluster, a pair of single-guide RNAs (sgRNAs) was designed to target the 5′ and 3′ flanking regions of the locus (Fig. S5A, B). The sgRNA sequences used were 5′-GCTGTTTATAACCGGGGCTT-3′ and 5′-GAAGGAACCTGCCGTCGCTC-3′, which were cloned into a pSpCas9(BB)-2A-puro (PX459) V2.0 vector (Addgene plasmid #62988; RRID: Addgene_62988; deposited by Feng Zhang). *DSP-*KO MDCK cells were generated using the CRISPR/Cas9 system with homology-independent DNA repair as described previously (Katoh et al., 2017). Briefly, a sgRNA targeting canine *DSP* (5′-GGTAATTTCCATCTGATCAT-3′) was cloned into pHiFiCas9 (Addgene plasmid #162277; Fujisawa et al., 2021) and pDonor-tBFP-NLS-Neo (Universal) (Addgene plasmid #80767; Katoh et al., 2017).

MDCK cells were transfected with the corresponding genome-editing constructs using Lipofectamine LTX and Plus Reagent (Life Technologies). For generation of *KRT-* null cells, cells were selected with 2 µg/mL puromycin for 48 h and cloned into 24-well plates. For generation of *DSP-*KO cells, cells were selected with 750 µg/mL G418 and cloned into 24-well plates. The loss of the keratin gene cluster was validated by genomic PCR and sequencing using specific primer sets (Fig. S5C and Supplementary Table S2), western blotting for KRT8 and KRT18, and pan-keratin immunofluorescence analysis. For *DSP-*KO cells, donor-vector integration and the genomic alterations were confirmed by genomic PCR and Sanger sequencing, respectively, using the primers listed in Supplementary Table S2. Loss of DSP protein expression was verified by immunoblotting and immunostaining (Fig. S2).

### Calcium-switch assay

Calcium-switch assays were performed under conditions similar to those described by Beggs et al. (2022). For synchronous junction assembly, confluent MDCK cells were incubated in calcium-free DMEM supplemented with 10% Chelex-treated FBS for at least 8 h. To prepare Chelex-treated FBS, FBS was mixed with Chele× 100 Resin (Bio-Rad, Hercules, CA, USA) and rotated overnight at 4°C, followed by filtration through a 0.22-µm filter. Junctional assembly was initiated by replacing the calcium-depleted medium with standard DMEM containing 1.8 mM Ca²⁺. The cells were fixed at 3 and 12 h after calcium restoration.

### TER measurement

Cells were cultured on Transwell filters (#3460; Corning Incorporated, Corning, NY, USA) for 4 days and equilibrated to room temperature before measurements. Resistance across the cell layer cultured on the filters was measured using an EVOM Manual (World Precision Instruments, Sarasota, FL, USA) equipped with STX4 electrodes (World Precision Instruments). Blank filter resistance was subtracted, and the filter area was multiplied to obtain the TER values (Ω·cm^2^).

### Western blotting

MDCK cells were washed with PBS and lysed in sodium dodecyl sulfate (SDS) sample buffer (62.5 mM Tris-HCl, pH 6.8, 5% 2-mercaptoethanol, 2% SDS, and 5% sucrose) to prepare whole-cell lysates. Cell lysates were separated using SDS– polyacrylamide gel electrophoresis and transferred to an Immobilon-P polyvinylidene difluoride membrane (Millipore, Burlington, MA, USA). The membranes were blocked with 5% non-fat skim milk in PBS containing 0.05% Tween-20 (PBS-T) and incubated with primary antibodies overnight at 4°C, followed by incubation with horseradish peroxidase (HRP)-conjugated secondary antibodies. Immunoreactive bands were detected using Immobilon Western Chemiluminescent HRP Substrate (Millipore) and ChemiDoc Touch Imaging System (Bio-Rad, Hercules, CA, USA). Uncropped images of the western blots are shown in Fig. S6.

### Image acquisition

For widefield fluorescence imaging, images were obtained using a THUNDER Ready microscope (Leica Microsystems, Wetzlar, Germany) equipped with HCX PL APO 100×/1.4–0.7 OIL or HCX PL APO 40×/1.25–0.75 OIL objectives and an ORCA-Flash4.0 Digital CMOS camera (HAMAMATSU Photonics, Hamamatsu City, Japan). Z-stack images were acquired at 0.2 µm intervals.

For apicobasal imaging of non-expanded samples, z-stacks were acquired using a ZEISS LSM 710 confocal microscope equipped with a Plan-Apochromat 63×/1.4 Oil DIC M27 objective with a free working distance of 0.19 mm for a 0.17-mm coverglass. ZEISS Immersol 518 F immersion oil (nₑ = 1.518 at 23°C) was used. Samples were imaged through high-precision #1.5H coverslips (0.170 ± 0.005 mm thickness) and mounted in Super Clear Mount (refractive index, 1.52). Z-stacks for apicobasal imaging were acquired at 0.44 µm intervals.

For apicobasal imaging of expanded samples, z-stacks were acquired using a ZEISS LSM 990 confocal microscope equipped with an LD C-Apochromat 40×/1.1 W Corr M27 water-immersion objective with a free working distance of 0.62 mm at a coverglass thickness of 0.17 mm and a correction-collar range of 0.15–0.19 mm. ZEISS Immersol W 2010 (nₑ = 1.334 at 23°C) was used as the immersion medium, and the correction collar was adjusted according to the thickness of the #1.5 glass bottom before imaging. Z-stacks for apicobasal imaging were acquired at 0.37 µm intervals. For super-resolution imaging of non-expanded samples in the calcium-switch analysis shown in Fig. S4I and J, images were acquired using a ZEISS LSM 990 equipped with an Airyscan 2 detector in SR mode and a Plan-Apochromat 63×/1.4 Oil DIC M27 objective (ZEISS). Airyscan images were reconstructed using joint deconvolution (jDCV) in ZEN software.

STED images were acquired using a Leica TCS SP8 STED 3X confocal microscope (Leica Microsystems) equipped with an HC PL APO CS2 100×/1.4 OIL objective, a white-light laser for excitation wavelengths between 470 and 670 nm, and a depletion laser at 660 nm. Images were acquired using LAS X software, followed by deconvolution using Huygens Pro software (Scientific Volume Imaging, Hilversum, The Netherlands).

### Image analysis

Image processing and distance measurements were performed using ImageJ software. Cell confluence was quantified as the percentage of the substrate area occupied by cells. Individual cell areas were measured by manually tracing cell boundaries defined by β-catenin staining. For the quantification of plaque-to-plaque distances, an analysis procedure based on previously established approaches for quantitative analysis of desmosomal nanoscale organization was used (Stahley et al., 2016; Beggs et al., 2022; Ainslie et al., 2025). Desmosomes were manually selected from single z-planes in which the two opposing DSP plaques were clearly resolved. Three independent one-pixel-wide line scans were drawn perpendicular to the cell–cell junction and across the opposing plaques at different positions within each desmosomal interface. For each line profile, the lowest fluorescence intensity value was defined as the local background and subtracted from the entire profile. The background-subtracted fluorescence intensity profiles were normalized and processed using a custom MATLAB-based pipeline incorporating Gaussian smoothing and spline interpolation. Peak positions corresponding to the opposing DSP plaques were identified using the peakfinder function with a consistent 40% intensity threshold, and the distance between the two peaks was calculated. The plaque-to-plaque distance for each desmosome was defined as the mean of the three measurements and plotted as a single data point. At least 30 individual desmosomes were analyzed for each condition.

The linear expansion factor was calculated as the ratio of the post-expansion gel diameter to the pre-expansion diameter (averaging 4.2 ± 0.1, n = 4 gels). To assess geometric preservation at the cellular scale, the same MDCK cells were imaged before and after expansion. Three vertices were selected within each cell to define two vectors (L₁ and L₂) originating from a common point and the angle between them (θ). Geometric preservation was assessed from the post-/pre-expansion ratio of the relative vector lengths (Rpost/Rpre, where R = L₁/L₂) and the included angle (θpost/θpre). At least 30 cells were analyzed in each of three independent experiments. At the desmosomal scale, we compared the DSP plaque-to-plaque distance and plaque length measured in non-expanded samples using STED microscopy with the corresponding dimensions measured in expanded samples using ExM. The effective lateral resolution of the ExM images was estimated to be approximately 60–70 nm, based on the approximately 4.2-fold expansion factor and a lateral optical resolution of approximately 250–300 nm under our imaging conditions. Plaque-to-plaque distances, image scale bars, and line-scan distance axes for the expanded samples are shown in post-expansion dimensions unless otherwise specified. For apicobasal analysis, fluorescence intensity profiles of OCLN, β-catenin, and DSG2 were generated along the z-axis using ImageJ. For each cell–cell interface, the position of maximum OCLN fluorescence intensity at each junction was defined as z = 0, and the positions of maximum DSG2 fluorescence intensities were determined relative to this reference. At least 30 independent cell–cell junctions were analyzed for each condition, with exact sample sizes provided in the corresponding figure legends. For fluorophore-swap analysis, the fluorophores assigned to OCLN and DSG2 were exchanged in both non-expanded and expanded samples, and their relative axial positions were quantified using the same procedure.

### Statistical analyses

For quantitative analyses of desmosomal organization performed across biological replicates, individual desmosomes were sampled from multiple fields of view within each biological replicate. Measurements from individual desmosomes were displayed to show the distribution of the data, together with mean ± s.d. Sample sizes and replication information are provided in the corresponding figure legends, as applicable. Statistical analyses were performed using Prism 10 software (GraphPad Software, San Diego, CA, USA). For comparisons between two groups, an unpaired Student’s *t*-test was used. For multiple-group comparisons, a one-way analysis of variance (ANOVA) followed by Tukey’s or Dunnett’s post hoc test, as appropriate, was performed. *P*-values are indicated in figures or figure legends, where statistical significance is defined as \**P* < 0.05, \*\**P* < 0.01, \*\*\**P* < 0.001, \*\*\*\**P* < 0.0001, and n.s. indicates not significant.

### Use of artificial intelligence tools

An OpenAI large language model was used during manuscript revision for consistency checking. The authors reviewed and edited the content as needed and take full responsibility for the content of the publication.

## Acknowledgments

We thank Kazuhisa Nakayama (Kumamoto University), Yohei Katoh (Hiroshima University), Mayumi Nakayama, and Yoko Furubayashi (Tohoku University) for technical assistance. The rat monoclonal anti-occludin antibody, MOC37, was kindly provided by Mikio Furuse (National Institute for Physiological Sciences) (Saitou et al., 1997). This work was supported by the Research Equipment Sharing System of Tohoku University (ZEISS LSM 990 with Airyscan 2), the MEXT Project for Promoting Public Utilization of Advanced Research Infrastructure (Program for Supporting Construction of Core Facilities; JPMXS0440600022), and Tohoku University Advanced Device Materials Platform under the MEXT Advanced Research Infrastructure for Materials and Nanotechnology in Japan (JPMXP1226TU0088).

## Footnotes

## Author contributions

Conceptualization: U.K. and S.C.; Data curation: U.K. and Y.A.; Formal analysis: U.K., Y.A. and T.O.; Funding acquisition: S.C., K.O. and U.K.; Investigation: U.K. and Y.A.; Methodology: S.C. and U.K.; Project administration: K.O. and S.C.; Resources: S.C.; Supervision: K.O. and S.C.; Validation: U.K., Y.A. and S.C.; Visualization: U.K., Y.A. and S.C.; Writing–original draft: U.K. and S.C.; Writing–review and editing: U.K., K.O., and S.C.

## Data and resource availability

All relevant data and resource details can be found in the article and its supplementary information. The MATLAB scripts used for plaque-to-plaque distance analysis are available from the corresponding author upon request.

## References

Ainslie, C. M., Patel, K., Tran, Y. T. B., Bartley, S. C., Bharathan, N. K., Spindler, V. and Mattheyses, A. L. (2025). The desmoplakin tail domain position in the desmosomal plaque is isoform dependent. J. Cell Sci. 138, jcs263906.

Barker, G. and Simmons, N. L. (1981). Identification of two strains of cultured canine renal epithelial cells (MDCK cells) which display entirely different physiological properties. Q. J. Exp. Physiol. 66, 61–72.

Beggs, R. R., Rao, T. C., Dean, W. F., Kowalczyk, A. P. and Mattheyses, A. L. (2022). Desmosomes undergo dynamic architectural changes during assembly and maturation. Tissue Barriers 10, 2017225.

Bharathan, N. K., Mattheyses, A. L. and Kowalczyk, A. P. (2024). The desmosome comes into focus. J. Cell Biol. 223, e202404120.

Chen, F., Tillberg, P. W. and Boyden, E. S. (2015). Expansion microscopy. Science 347, 543–548.

Coulombe, P. A. and Omary, M. B. (2002). ‘Hard’ and ‘soft’ principles defining the structure, function and regulation of keratin intermediate filaments. Curr. Opin. Cell Biol. 14, 110–122.

Cui, Y., Yang, G., Goodwin, D. R., O’Flanagan, C. H., Sinha, A., Zhang, C., Kitko, K. E., Shin, T. W., Park, D., Aparicio, S., et al. (2023). Expansion microscopy using a single anchor molecule for high-yield multiplexed imaging of proteins and RNAs. PLoS ONE 18, e0291506.

Dong, Y., Elgerbi, A., Xie, B., Han, Y., Kwiatkowski, A. V., Choy, J. S. and Sivasankar, S. (2025). Actomyosin forces trigger a conformational change in desmoplakin within desmosomes. Nat. Commun. 16, 9052.

Farquhar, M. G. and Palade, G. E. (1963). Junctional complexes in various epithelia. J. Cell Biol. 17, 375–412.

Fujisawa, S., Qiu, H., Nozaki, S., Chiba, S., Katoh, Y. and Nakayama, K. (2021). ARL3 and ARL13B GTPases participate in distinct steps of INPP5E targeting to the ciliary membrane. Biol. Open 10, bio058843.

Furuse, M., Furuse, K., Sasaki, H. and Tsukita, S. (2001). Conversion of zonulae occludentes from tight to leaky strand type by introducing claudin-2 into Madin-Darby canine kidney I cells. J. Cell Biol. 153, 263–272.

Hagiyama, M., Yabuta, N., Okuzaki, D., Inoue, T., Takashima, Y., Kimura, R., Ri, A. and Ito, A. (2017). Modest static pressure suppresses columnar epithelial cell growth in association with cell shape and cytoskeletal modifications. Front. Physiol. 8, 997.

Harmon, R. M. and Green, K. J. (2013). Structural and functional diversity of desmosomes. Cell Commun. Adhes. 20, 171–187.

Herrmann, H. and Aebi, U. (2016). Intermediate filaments: structure and assembly. Cold Spring Harb. Perspect. Biol. 8, a018242.

Janssen, V. and Huveneers, S. (2024). Cell–cell junctions in focus – imaging junctional architectures and dynamics at high resolution. J. Cell Sci. 137, jcs262041.

Katoh, Y., Michisaka, S., Nozaki, S., Funabashi, T., Hirano, T., Takei, R. and Nakayama, K. (2017). Practical method for targeted disruption of cilia-related genes by using CRISPR/Cas9-mediated, homology-independent knock-in system. Mol. Biol. Cell 28, 898–906.

Moch, M., Schwarz, N., Windoffer, R. and Leube, R. E. (2020). The keratin– desmosome scaffold: pivotal role of desmosomes for keratin network morphogenesis. Cell. Mol. Life Sci. 77, 543–558.

Moch, M., Schieren, J. and Leube, R. E. (2022). Cortical tension regulates desmosomal morphogenesis. Front. Cell Dev. Biol. 10, 946190.

Ninomiya, K., Ohta, K., Kawasaki, U., Chiba, S., Inoue, T., Kuranaga, E., Ohashi, K. and Mizuno, K. (2024). Calcium influx promotes PLEKHG4B localization to cell–cell junctions and regulates the integrity of junctional actin filaments. Mol. Biol. Cell 35, ar24.

North, A. J., Bardsley, W. G., Hyam, J., Bornslaeger, E. A., Cordingley, H. C., Trinnaman, B., Hatzfeld, M., Green, K. J., Magee, A. I. and Garrod, D. R. (1999). Molecular map of the desmosomal plaque. J. Cell Sci. 112, 4325–4336.

Rajasekaran, A. K., Hojo, M., Huima, T. and Rodriguez-Boulan, E. (1996). Catenins and zonula occludens-1 form a complex during early stages in the assembly of tight junctions. J. Cell Biol. 132, 451–463.

Saitou, M., Ando-Akatsuka, Y., Itoh, M., Furuse, M., Inazawa, J., Fujimoto, K. and Tsukita, S. (1997). Mammalian occludin in epithelial cells: its expression and subcellular distribution. Eur. J. Cell Biol. 73, 222–231.

Schmitt, T., Huber, J., Pircher, J., Schmidt, E. and Waschke, J. (2025). The impact of signaling pathways on the desmosome ultrastructure in pemphigus. Front. Immunol. 15, 1497241.

Stahley, S. N., Bartle, E. I., Atkinson, C. E., Kowalczyk, A. P. and Mattheyses, A. L. (2016). Molecular organization of the desmosome as revealed by direct stochastic optical reconstruction microscopy. J. Cell Sci. 129, 2897–2904.

Stappenbeck, T. S., Bornslaeger, E. A., Corcoran, C. M., Luu, H. H., Virata, M. L. and Green, K. J. (1993). Functional analysis of desmoplakin domains: specification of the interaction with keratin versus vimentin intermediate filament networks. J. Cell Biol. 123, 691–705.

Tillberg, P. W., Chen, F., Piatkevich, K. D., Zhao, Y., Yu, C.-C., English, B. P., Gao, L., Martorell, A., Suk, H.-J., Yoshida, F., et al. (2016). Protein-retention expansion microscopy of cells and tissues labeled using standard fluorescent proteins and antibodies. Nat. Biotechnol. 34, 987–992.

Tokuda, S. and Furuse, M. (2015). Claudin-2 knockout by TALEN-mediated gene targeting in MDCK cells: claudin-2 independently determines the leaky property of tight junctions in MDCK cells. PLoS ONE 10, e0119869.

Vijayaraj, P., Kröger, C., Reuter, U., Windoffer, R., Leube, R. E. and Magin, T. M. (2009). Keratins regulate protein biosynthesis through localization of GLUT1 and -3 upstream of AMP kinase and Raptor. J. Cell Biol. 187, 175–184.

Yeruva, S. and Waschke, J. (2023). Structure and regulation of desmosomes in intercalated discs: lessons from epithelia. J. Anat. 242, 81–90.

