## Supplementary Figures and Tables for "Density-Dependent Rearrangement of Desmoplakin in Epithelial Junctions: Insights from Expansion Microscopy"

**Figure S1. Validation of ExM expansion and desmosomal measurements.**

(A, B) Macroscopic expansion of the hydrogel. (A) Representative images of the hydrogel before and after expansion. (B) Post-/pre-expansion gel diameter ratio (n = 4 gels). (C, D) Same-cell analysis before and after expansion: representative DSP images (C) and post-/pre-expansion ratios of relative vector lengths (Rpost/Rpre, where $R = L_1/L_2$) and included angles ($\theta$post/$\theta$pre) (D). At least 30 cells were analyzed in each of three independent experiments. (E–H) Comparison of desmosomal dimensions measured by ExM and STED. (E) Representative images of DSP C-terminus obtained by ExM and STED. (F, G) DSP C-terminal plaque-to-plaque distance and plaque length measured by ExM (F) and STED (G). (H) Ratios of individual ExM measurements to the corresponding mean STED values. For F and G, 44 and 48 desmosomes were analyzed by ExM and STED, respectively. Data represent mean ± s.d. Scale bars: 10 μm (pre-expansion) and 40 μm (post-expansion) (C); 20, 2 and 0.8 μm (ExM) and 5, 0.5 and 0.2 μm (STED), from overview to successive magnifications (E).

**Figure S2. Functional validation of mSca3-DSP in *DSP*-KO MDCK cells.**

(A) Immunoblot analysis of DSP in WT and *DSP*-KO MDCK cells. β-actin was used as a loading control. (B) Indels at the *DSP* target site in *DSP*-KO cells were confirmed by direct sequencing following genomic PCR verification of donor-vector integration (Katoh et al., 2017). (C, D) *DSP*-KO MDCK cells expressing mSca3-DSP: representative ExM images stained for DSP C-terminus and mCherry (DSP N-terminus) (C), with corresponding line-scan profiles along the dotted line (D). (E, F) Representative images of *DSP*-KO cells with or without mSca3-DSP expression, stained for mSca3, DSG2, and DSP (E), or mSca3, pan-keratin (KRT), and PKP2 (F). For *DSP*-KO cells, the same magnified regions are also shown with saturated display settings to visualize residual junctional signals. In *DSP*-KO cells, junctional accumulation of DSG2 and PKP2 and the association of keratin filaments with desmosomal junctions were reduced; these features were restored by mSca3-DSP expression. Scale bars: 20, 4 and 1 μm (C), from overview to successive magnifications; 10 and 1 μm (E, F), from overview to magnification.

**Figure S3. Validation of cell-density conditions and fluorophore-swap analysis.** (A–C) Characterization of subconfluent (Sub.) and confluent (Con.) MDCK cultures. (A) Representative low-magnification images stained for β-catenin, F-actin, and nuclei. (B, C) Quantification of cell area (B) and cell confluence (substrate occupancy; C). Cell area analysis included 45–55 cells per condition per replicate, and cell confluence analysis included five fields per condition per replicate. Confluent cultures showed greater substrate occupancy and smaller cell areas than subconfluent cultures, confirming distinct epithelial density states. (D, E) Fluorophore-swap analysis of non-expanded confluent MDCK cells stained for DSG2, OCLN, and β-catenin (β-cat), showing an XZ view (D) and relative fluorescence intensity profile along the z-axis (E). (F, G) Corresponding analysis of expanded confluent cells, showing an XZ view (F) and relative fluorescence intensity profile along the z-axis (G). Mean profiles in E and G were averaged from 60 and 43 independent cell–cell junctions, respectively; error bars indicate s.d. DSG2 remained basal to OCLN after fluorophore exchange in both non-expanded and expanded samples. Imaging conditions used to minimize refractive-index mismatch and associated spherical aberration are described in the Materials and Methods. Scale bars: 50 µm (A), 1 µm (D, F).

**Figure S4. Validation of DSP plaque-spacing measurements under different experimental conditions.**

(A, B) ExM analysis of subconfluent (Sub.) and confluent (Con.) MDCK cells fixed with methanol (MeOH; −20°C) or 50% methanol/50% acetone (MeOH/acetone; 4°C): representative images of DSP C-terminus (A) and plaque-to-plaque distance (B). (C, D) ExM analysis of subconfluent and confluent MDCK cells co-stained with anti-DSP antibodies 651109 and A303-356A: representative images (C) and DSP C-terminal plaque-to-plaque distance (D). Under both fixation conditions and with both DSP antibodies, DSP C-terminal spacing was greater in confluent than subconfluent cells. (E, F) Non-expanded subconfluent and confluent MDCK cells analyzed by STED: representative images stained for DSP C-terminus and DSG2 (E), and DSP C-terminal plaque-to-plaque distance (F). STED analysis of non-expanded samples also showed greater DSP C-terminal spacing in confluent than subconfluent cells. (G, H) Calcium-switch analysis at 3 and 12 h after calcium restoration: representative ExM images of DSP C-terminus (G) and plaque-to-plaque distance (H). (I, J) Corresponding analysis of non-expanded samples at 3 and 12 h after calcium restoration using a ZEISS LSM 990 in SR mode with a Plan-Apochromat 63×/1.4 Oil objective: representative images (I) and DSP C-terminal plaque-to-plaque distance (J). DSP C-terminal spacing was greater at 12 h than at 3 h after calcium restoration in both ExM and non-expanded SR analyses. (K) TER of confluent MDCK monolayers. TER was $113 \pm 11$ $\Omega \cdot cm^2$ ($n = 3$ independent biological replicates). For the ExM analyses in B, D and H, plaque-to-plaque distances are shown in post-expansion dimensions. For B, D, F, H and J, each dot represents one desmosome. For B, 50, 50, 34 and 33 desmosomes were analyzed for the four conditions, respectively; for D, 43, 43, 40 and 40 desmosomes were analyzed for the four conditions, respectively; for F, 52 and 55 desmosomes were analyzed under subconfluent and confluent conditions, respectively; for H, 72 and 70 desmosomes were analyzed at 3 and 12 h, respectively; for J, 36 and 40 desmosomes were analyzed at 3 and 12 h, respectively. Horizontal lines and error bars indicate the mean and s.d., respectively. Scale bars: 20, 4 and 1 µm (A, C), from overview to successive magnifications; 0.5 and 0.2 µm (E, I); 2 and 0.8 µm (G), from overview to magnification.

**Figure S5. Generation and validation of *KRT*-null MDCK cells and model of DSP reorganization.**

(A, B) Schematic of CRISPR/Cas9-mediated deletion of the type-II keratin gene cluster using two sgRNAs targeting its 5′ and 3′ ends. (C) Genomic PCR strategy and validation of the deletion. Primer sets A and B amplified WT genomic DNA, whereas primer set C amplified only *KRT*-null genomic DNA. (D) Representative images of WT and *KRT*-null MDCK cells stained for pan-keratin (KRT), DSP, and DAPI. (E) Immunoblot analysis of pan-keratin, KRT8, and KRT18 in WT and *KRT*-null MDCK cells. Increasing amounts of WT cell lysate were loaded as indicated by the triangle. α-tubulin was used as a loading control. *KRT*-null cells showed loss of keratin protein expression and filamentous pan-keratin staining. (F) Schematic summary of density-associated DSP plaque organization in WT and *KRT*-null MDCK cells. In WT cells, plaque-to-plaque spacing increases from subconfluent to confluent conditions for both the DSP N- and C-terminal signals, with a larger increase at the C-terminus. In *KRT*-null cells, DSP C-terminal spacing is already elevated under subconfluent conditions and shows no further density-associated increase. Magenta indicates DSP, and green indicates keratin filaments; pale green traces in *KRT*-null cells indicate the absence of keratin filament formation. The schematic summarizes

739 the relative positions of DSP N- and C-terminal signals and the keratin filament network

740 and is not drawn to scale. Scale bar: 15 µm (D).

741

742 **Figure S6. Uncropped western blot images.**

743 Uncropped western blot images corresponding to the immunoblots shown in Fig. S2A

744 and Fig. S5E.

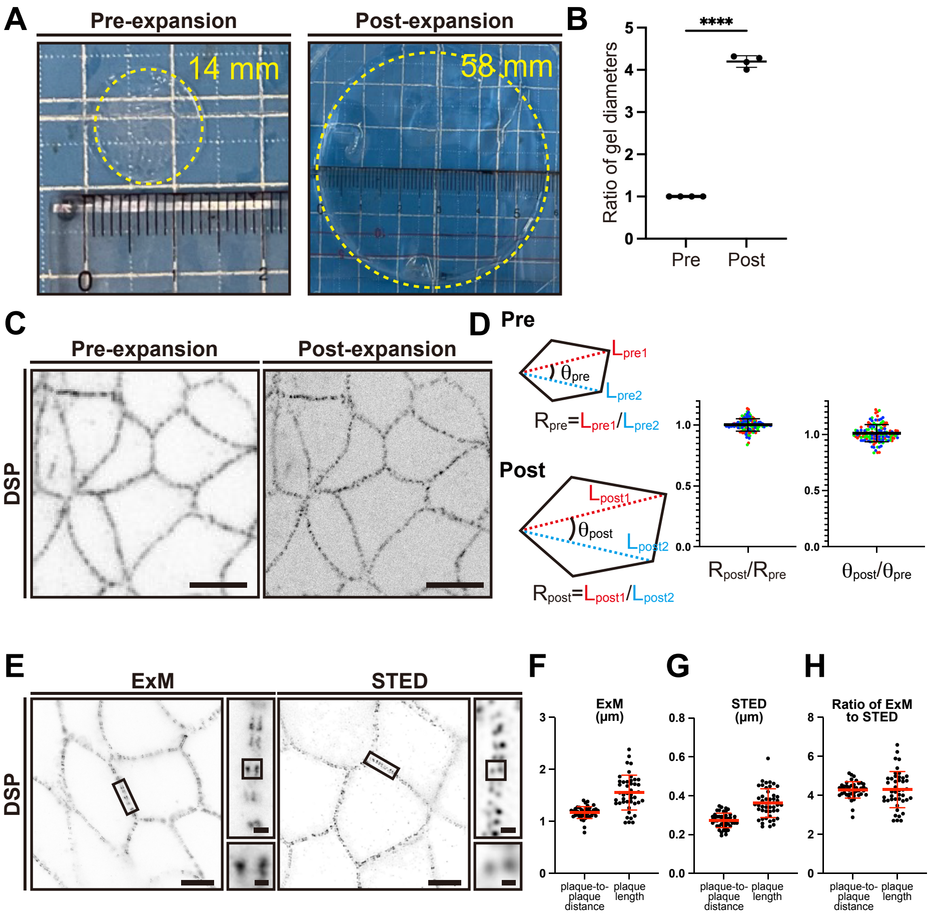
A
Pre-expansion
Post-expansion
14 mm
58 mm
B
****
Ratio of gel diameters
Pre
Post
C
Pre-expansion
Post-expansion
DSP
D
Pre
Post
$R_{pre}=L_{pre1}/L_{pre2}$
$R_{post}=L_{post1}/L_{post2}$
$R_{post}/R_{pre}$
$\theta_{post}/\theta_{pre}$
E
ExM
STED
DSP
F
ExM (μm)
G
STED (μm)
H
Ratio of ExM to STED
plaque-to-plaque distance
plaque length

**Figure S1**

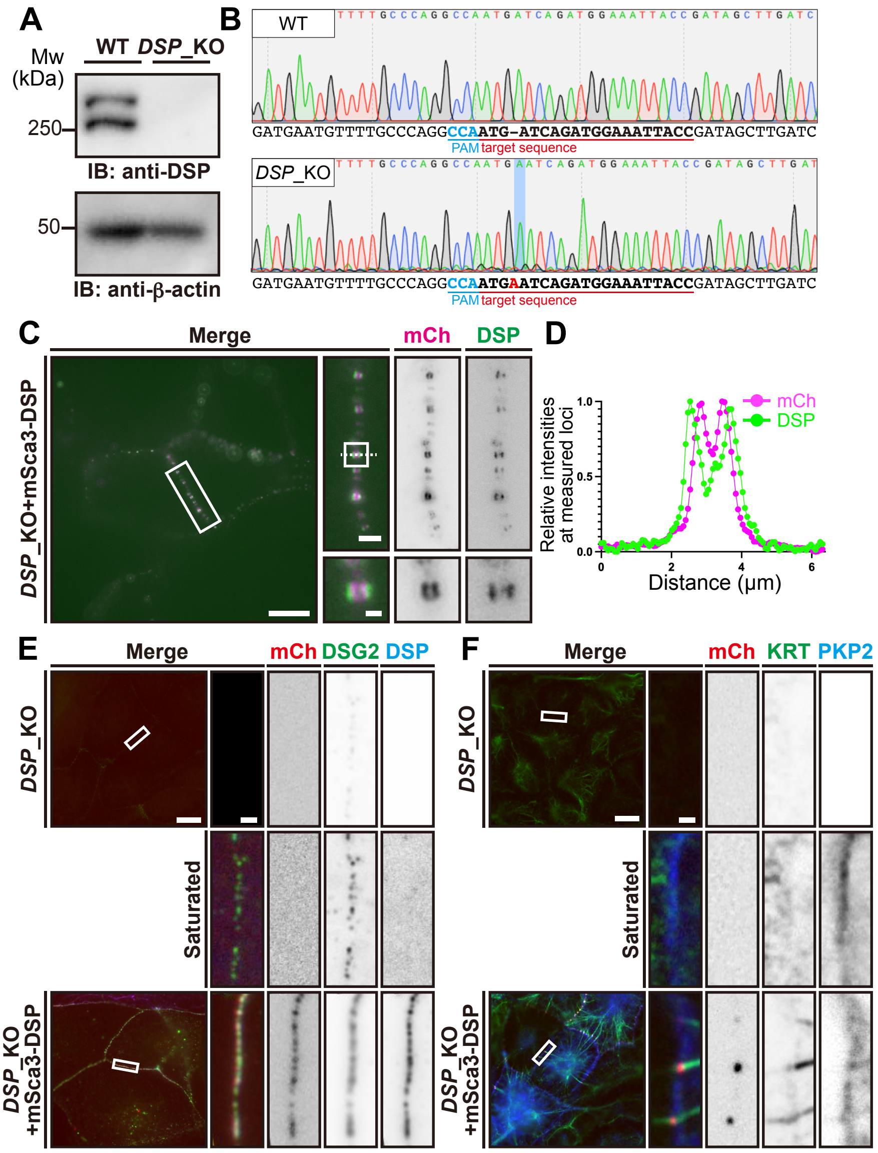

A
Mw (kDa)
WT DSP_KO
250
IB: anti-DSP
50
IB: anti-β-actin
B
WT
GATGAATGTTTTGCCCAGGCCAATG-ATCAGATGGAAATTACCGATAGCTTGATC
PAM target sequence
DSP_KO
GATGAATGTTTTGCCCAGGCCAATGAATCAGATGGAAATTACCGATAGCTTGATC
PAM target sequence
C
Merge
mCh
DSP
DSP_KO+mSca3-DSP
D
Relative intensities at measured loci
mCh
DSP
Distance (μm)
E
Merge
mCh DSG2 DSP
DSP_KO
Saturated
DSP_KO +mSca3-DSP
F
Merge
mCh KRT PKP2
DSP_KO
Saturated
DSP_KO +mSca3-DSP


**Figure S2**

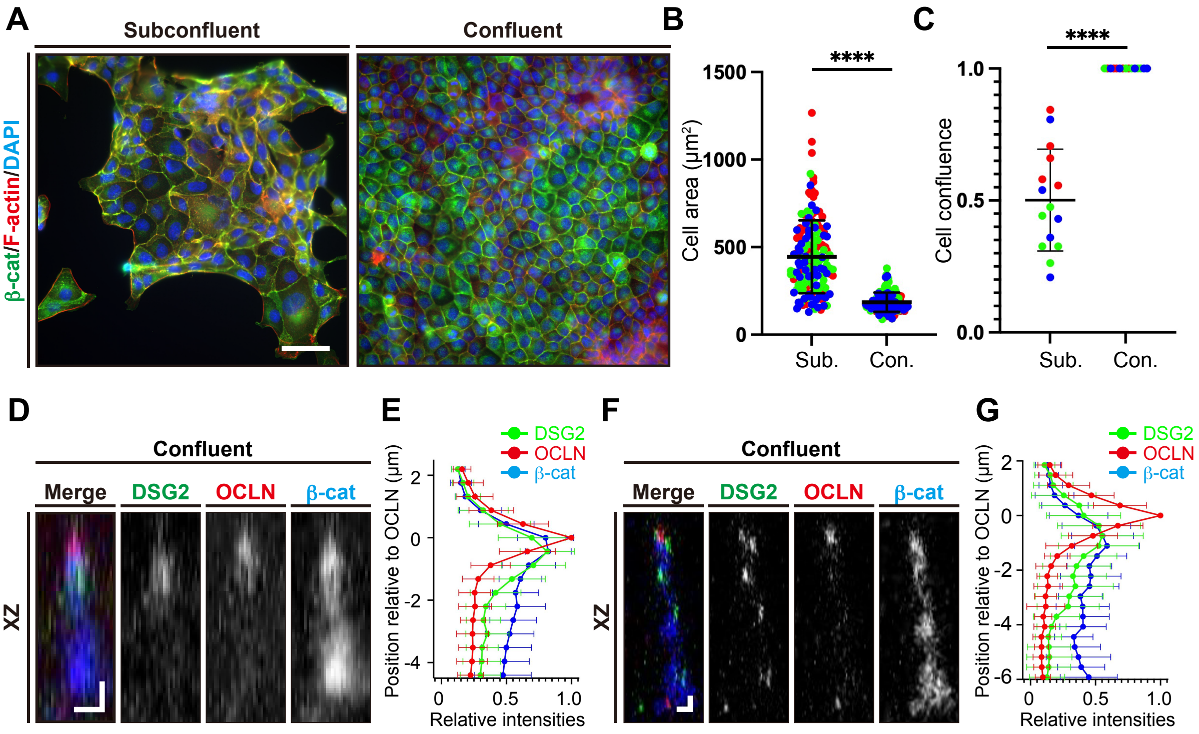

A
Subconfluent
Confluent
β-cat/F-actin/DAPI
B
****
1500
1000
500
0
Cell area (μm²)
Sub.
Con.
C
****
1.0
0.5
0.0
Cell confluence
Sub.
Con.
D
Confluent
Merge
DSG2
OCLN
β-cat
XZ
E
DSG2
OCLN
β-cat
Position relative to OCLN (μm)
Relative intensities
F
Confluent
Merge
DSG2
OCLN
β-cat
XZ
G
DSG2
OCLN
β-cat
Position relative to OCLN (μm)
Relative intensities


**Figure S3**

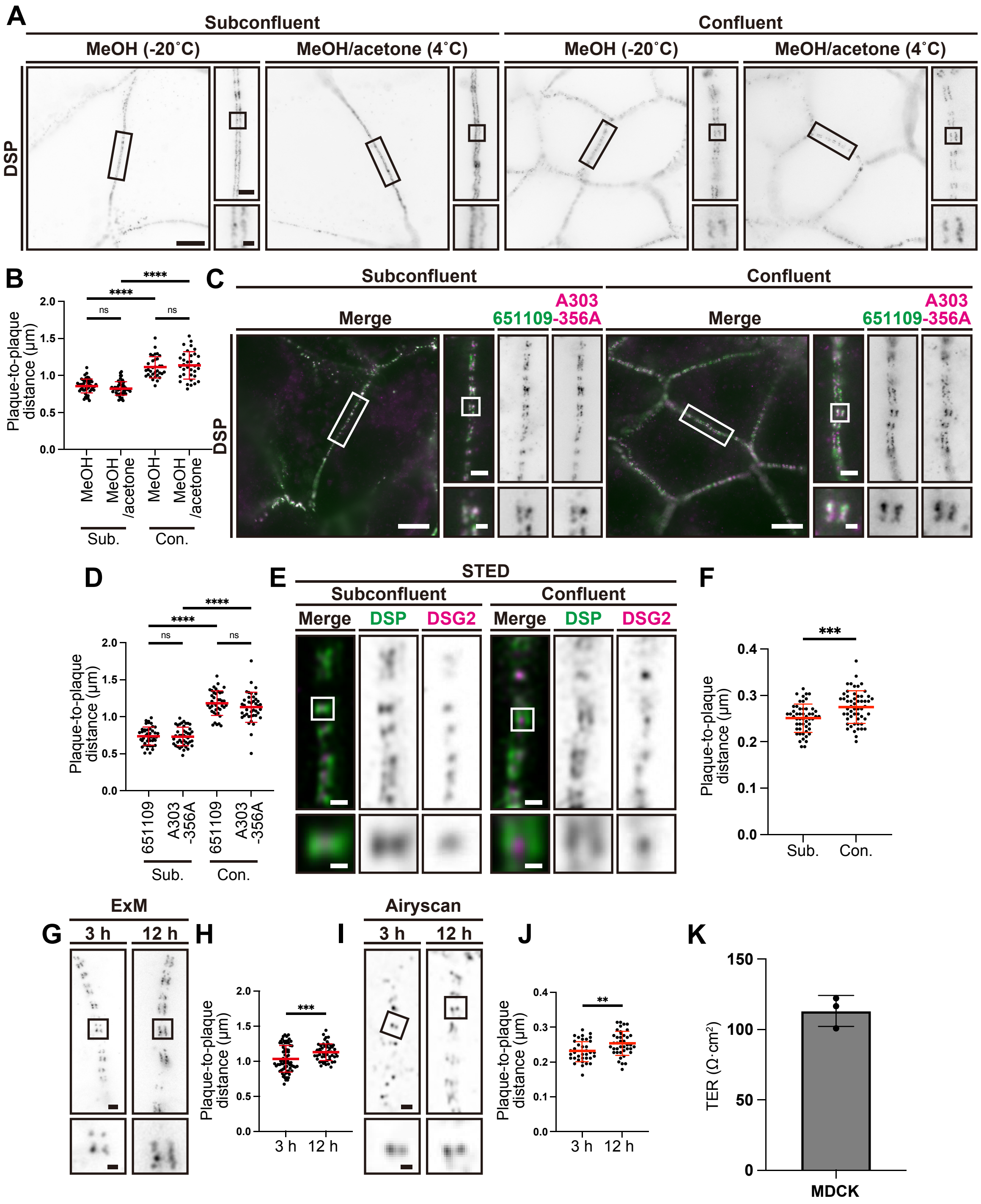
A
Subconfluent
Confluent
MeOH (-20˚C)
MeOH/acetone (4˚C)
DSP
B
Plaque-to-plaque distance (μm)
MeOH
MeOH /acetone
Sub.
Con.
C
Merge
651109-A303-356A
D
651109
A303 -356A
E
STED
DSP
DSG2
F
G
ExM
3 h
12 h
H
I
Airyscan
J
K
TER (Ω·cm²)
MDCK

**Figure S4**

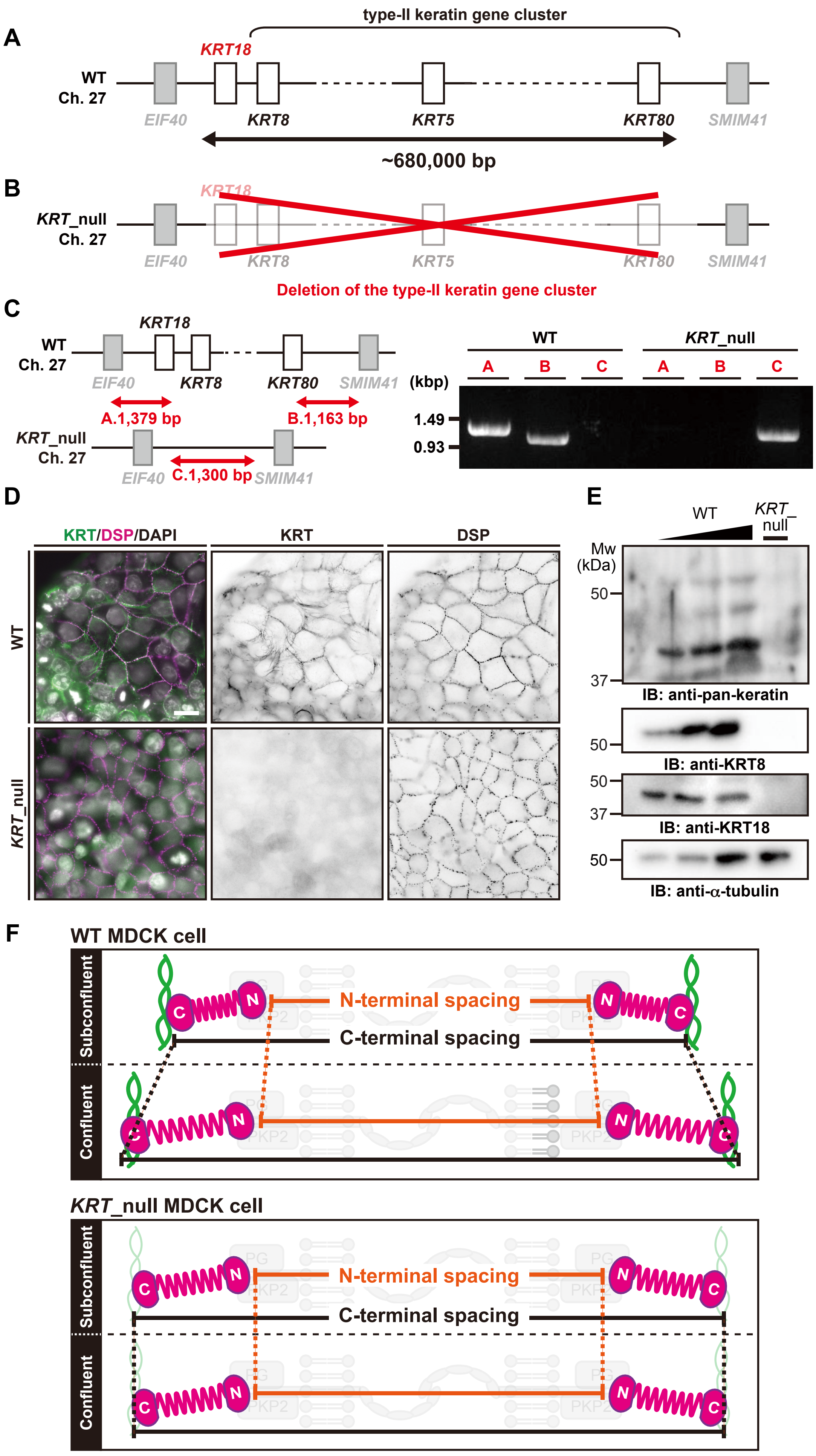
A
type-II keratin gene cluster
KRT18
WT
Ch. 27
EIF40
KRT8
KRT5
KRT80
SMIM41
~680,000 bp
B
KRT_null
Ch. 27
Deletion of the type-II keratin gene cluster
C
A.1,379 bp
B.1,163 bp
C.1,300 bp
(kbp)
1.49
0.93
D
KRT/DSP/DAPI
KRT
DSP
E
Mw (kDa)
50
37
IB: anti-pan-keratin
IB: anti-KRT8
IB: anti-KRT18
IB: anti-α-tubulin
F
WT MDCK cell
Subconfluent
Confluent
N-terminal spacing
C-terminal spacing
KRT_null MDCK cell
PG
PKP2

Figure S5

Figure S2

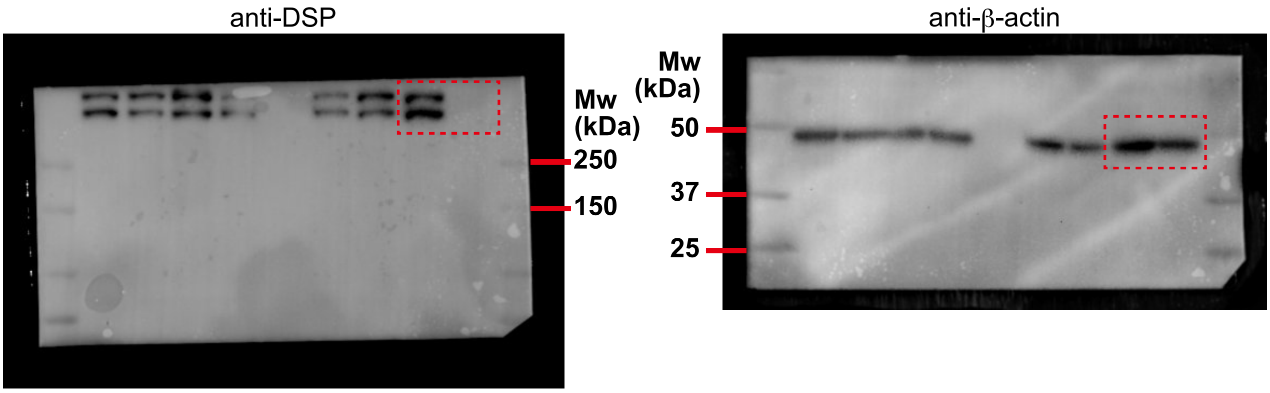


Figure S5

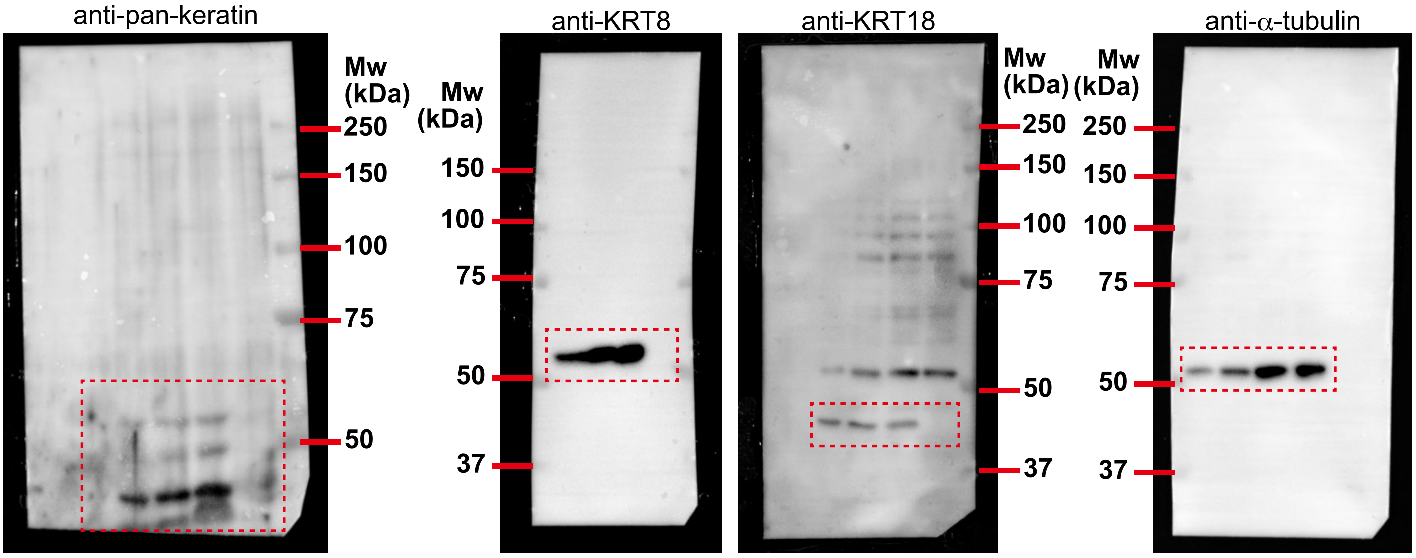


**Figure S6**

Table S1. Antibodies used in this study

| Antibody | Manufacturer or provider | Clone/catalog number or reference number | Dilution (purpose) |
|---|---|---|---|
| Monoclonal-anti-Desmoplakin | Progen | 651109 | 1:500 (IF; nonEx), 1:100 (IF; Ex) |
| Polyclonal-anti-Desmoplakin | Bethyl Lab. | A303-356A | 1:250 (IF; Ex) |
| Monoclonal-anti-β-catenin | BD Biosciences | 610154 | 1:500 (IF; nonEx), 1:250 (IF; Ex) |
| Monoclonal-anti-plakoglobin | BD Biosciences | 610253 | 1:500 (IF; nonEx), 1:250 (IF; Ex) |
| Polyclonal-anti-wide spectrum Cytokeratin | Abcam | ab9377 | 1:500 (IF; nonEx), 1:250 (IF; Ex), 1:1,000 (IB) |
| Polyclonal-anti-Desmoglein 2 | Proteintech | 21880-1-AP | 1:500 (IF; nonEx), 1:250 (IF; Ex) |
| Monoclonal-anti-plakophilin 2 | Progen | 651101 | 1:500 (IF; nonEx), 1:250 (IF; Ex) |
| Monoclonal-anti-Occludin | Mikio Furuse, National Institute for Physiological Sciences (gift) | MOC37 | 1:100 (IF; nonEx), 1:50 (IF; Ex) |
| Polyclonal-anti-mCherry | Proteintech | 26765-1-AP | 1:500 (IF; nonEx), 1:250 (IF; Ex) |
| Monoclonal-anti-Keratin 8 | Sigma-Aldrich | MABT329 | 1:1,000 (IB) |
| Monoclonal-anti-Keratin 18 | Progen | 61028 | 1:1,000 (IB) |
| Monoclonal-anti-β-actin | Abcam | ab6276 | 1:1,500 (IB) |
| Monoclonal-anti-α-tubulin | Abcam | B-5-1-2 | 1:2,000 (IB) |
| Alexa Fluor 488-conjugated goat anti-mouse IgG | Invitrogen | A11029 | 1:500 (IF; nonEx), 1:250 (IF; Ex) |
| Alexa Fluor 555-conjugated goat anti-mouse IgG | Invitrogen | A31570 | 1:300 (IF; nonEx) |
| Alexa Fluor 568-conjugated goat anti-mouse IgG (H+L) | Invitrogen | A11031 | 1:500 (IF; nonEx), 1:250 (IF; Ex) |
| Alexa Fluor 633-conjugated goat anti-mouse IgG (H+L) | Invitrogen | A21052 | 1:500 (IF; nonEx), 1:250 (IF; Ex) |
| Alexa Fluor 488-conjugated goat anti-mouse IgG2a | Invitrogen | A21131 | 1:500 (IF; nonEx), 1:250 (IF; Ex) |
| Alexa Fluor 568-conjugated goat anti-mouse IgG2b | Invitrogen | A21144 | 1:500 (IF; nonEx), 1:100 (IF; Ex) |
| Alexa Fluor 488-conjugated goat anti-rabbit IgG (H+L) | Invitrogen | A11034 | 1:500 (IF; nonEx), 1:250 (IF; Ex) |

| | | | |
|---|---|---|---|
| Alexa Fluor 568-conjugated goat anti-rabbit IgG (H+L) | Invitrogen | A11011 | 1:500 (IF; nonEx), 1:250 (IF; Ex) |
| Alexa Fluor 488-conjugated donkey anti-rat IgG (H+L) | Invitrogen | A11006 | 1:500 (IF; nonEx), 1:250 (IF; Ex) |
| Alexa Fluor 568-conjugated donkey anti-rat IgG (H+L) | Invitrogen | A78946 | 1:500 (IF; nonEx), 1:250 (IF; Ex) |
| CF568 donkey anti-rabbit IgG (H+L) | Biotium | 20098 | 1:1,000 (IF; nonEx), 1:500 (IF; Ex) |
| CF633 donkey anti-mouse IgG (H+L) | Biotium | 20124 | 1:1,000 (IF; nonEx), 1:500 (IF; Ex) |
| CF633 goat anti-mouse IgG1(γ1) | Biotium | 20250 | 1:1,000 (IF; nonEx), 1:500 (IF; Ex) |
| Rhodamine phalloidin | FUJIFILM | 181-02921 | 1:500 (IF; nonEx) |
| DAPI | DOJINDO | 340-07971 | 20 ng/mL (IF; nonEx), 40 ng/mL (IF; Ex) |
| Horseradish peroxidase (HRP)-conjugated sheep anti-mouse IgG | Sigma-Aldrich | NA931 | 1:10,000 (IB) |
| Horseradish peroxidase (HRP)-conjugated donkey anti-rabbit IgG | Sigma-Aldrich | NA934 | 1:10,000 (IB) |

IF; immunofluorescence; IB, immunoblotting; nonEx, non-expanded sample; Ex; expanded sample

Table S2. Primers used for genomic PCR in this study

| Name | Sequence |
|---|---|
| KRT_deletion_genomicPCR_Fw | 5′-GTCCAATTTAGGAAATTATTTTACAGTGGT-3′ |
| KRT_control_genomicPCR_Rv | 5′-CTGCTAAAACTTGCTGAGACTATCCTT-3′ |
| KRT_control_genomicPCR_Fw | 5′-TGCTTTTTCTCTGCCTATTTATTGG-3′ |
| KRT_deletion_genomicPCR_Rv | 5′-CTTGAAACCTTGAAACAAAAGGTGT-3′ |
| DSP_KO_genomicPCR_Fw | 5′-TTCTTTGTAAATATGAAAGATAGCGTTTAACAC-3′ |
| DSP_KO_genomicPCR_Rv | 5′-TGCTCTCTGACTCTACAAGTCTCATTTTTATAT-3′ |
| pDonor_tagBFP_Rv | 5′-GCACTTGAAGTGATGGTTGTC-3′ |
